# The Neutrophil-to-Lymphocyte Ratio as a Prognostic Indicator in Head and Neck Cancer: A Systematic Review and Meta-Analysis

**DOI:** 10.1101/217034

**Authors:** Tristan Tham, Yonatan Bardash, Saori Wendy Herman, Peter D. Costantino

**Affiliations:** New York Head & Neck Institute, Hofstra Northwell School of Medicine, Northwell Health System, New York, USA; Hofstra Northwell School of Medicine, New York, USA

**Keywords:** Neutrophil Lymphocyte Ratio, NLR, Inflammatory Markers, Prognosis, Head and Neck Cancer, Meta-Analysis, Systematic Review

## Abstract

**Background:** The aim of this systematic review and meta-analysis was to investigate the relationship between the Neutrophil-to-Lymphocyte Ratio (NLR) and prognosis in HNC.

**Methods:** Studies were identified from Pubmed, Embase, Scopus, and the Cochrane Library. A systematic review and meta-analysis were performed to generate the pooled hazard ratios (HR) for overall survival (OS), disease free survival (DFS), and progression free survival (PFS).

**Results:** Our analysis combined the results of over 6770 patients in 26 cohorts (25 studies). The pooled data demonstrated that an elevated NLR significantly predicted poorer OS, DFS, and PFS. Heterogeneity was found for OS, PFS, and marginally for DFS. Subgroup analysis in OS demonstrated that elevated NLR remained an indicator of poor prognosis.

**Conclusions:** Elevated pretreatment NLR is a prognostic marker for HNC. It represents a simple and easily obtained marker that could be used to stratify groups of high-risk patients that might benefit from adjuvant therapy.

## Introduction

Head and neck cancer (HNC) is one of the more common cancers worldwide, ^1^ accounting for more than half a million new cases annually. The majority of HNCs are of the squamous cell carcinoma (SCC) histological subtype, and may be located in the anatomical compartments of the oral cavity, nasopharynx, oropharynx, hypopharynx, and larynx. Standard therapies of HNC may be surgical, radiotherapy or chemotherapy, or a combination thereof. The mode of treatment is largely determined by the characteristics of the presenting tumor, namely the stage, grade, and location. These in turn determine the prognosis of the tumor. The other known prognostic factors for HNC include performance status, smoking and alcohol history, and human papillomavirus (HPV) infection.

Recently, there has been an interest in easily obtained inflammatory biomarkers that have the potential to predict the prognosis in patients with cancer. Such markers are hypothesized to reflect the underlying complex interplay between the systemic inflammatory responses with the tumor microenvironment ^2–4^. Markers such as CRP, neutrophil-to-lymphocyte ratio (NLR), platelet-to-lymphocyte ratio (PLR), lymphocyte-to-monocyte ratio (LMR) have been described in the literature. Of these inflammatory markers, the NLR has been widely reported. The ostensible ability of the NLR to act as a prognostic tool has been demonstrated in several meta-analyses in different cancer sites ^5–7^.

Despite the surfeit amount of studies published, the prognostic value of the NLR in HNC patients remains unclear, and even controversial. ^8^ There are several meta-analyses of the NLR in nasopharyngeal cancers only ^9, 10^. Therefore, our aim in this study was to consolidate the published literature, in order to clarify the relationship between the pretreatment NLR and the prognosis of patients with cancer in all sites of the head and neck. To the best of our knowledge, this is the first meta-analysis investigating the prognostic role of NLR in all sites of the head and neck.

## Materials and Methods

### Design

Our search was performed in accordance with the Cochrane Handbook of DTA Chapter on searching ^11^. We followed the Preferred Reporting Items for Systematic Reviews and Meta-Analyses (PRISMA) statement guidelines to identify, screen, and describe the protocols used in this systematic review. ^12^ Since our systematic review and meta-analysis were performed on observational studies, we also followed the Meta-analysis Of Observational Studies in Epidemiology (MOOSE) Checklist. ^13^ Our search strategy was designed in collaboration with a librarian at the Hofstra Northwell School of Medicine (WH), and the systematic review was prospectively registered in an online systematic review database (PROSPERO 2017:CRD42017059500).^14^

### Search Strategy

PubMed (via the web), EMBASE, and the Cochrane Library were searched on March 17, 2017. Scopus was searched on March 20, 2017. We searched all databases from their inception to the present with no restriction on language of publication. To gather additional literature, bibliographies were hand searched and PubMed’s related articles search was performed on all included articles. Due to the large volume of results retrieved in Embase and Scopus, the publication type filters were used to exclude conference abstracts, letters, editorials, conference reviews, conference papers, and book chapters. A detailed description of the search strategy and results may be found in Table 1.

**Table 1.**
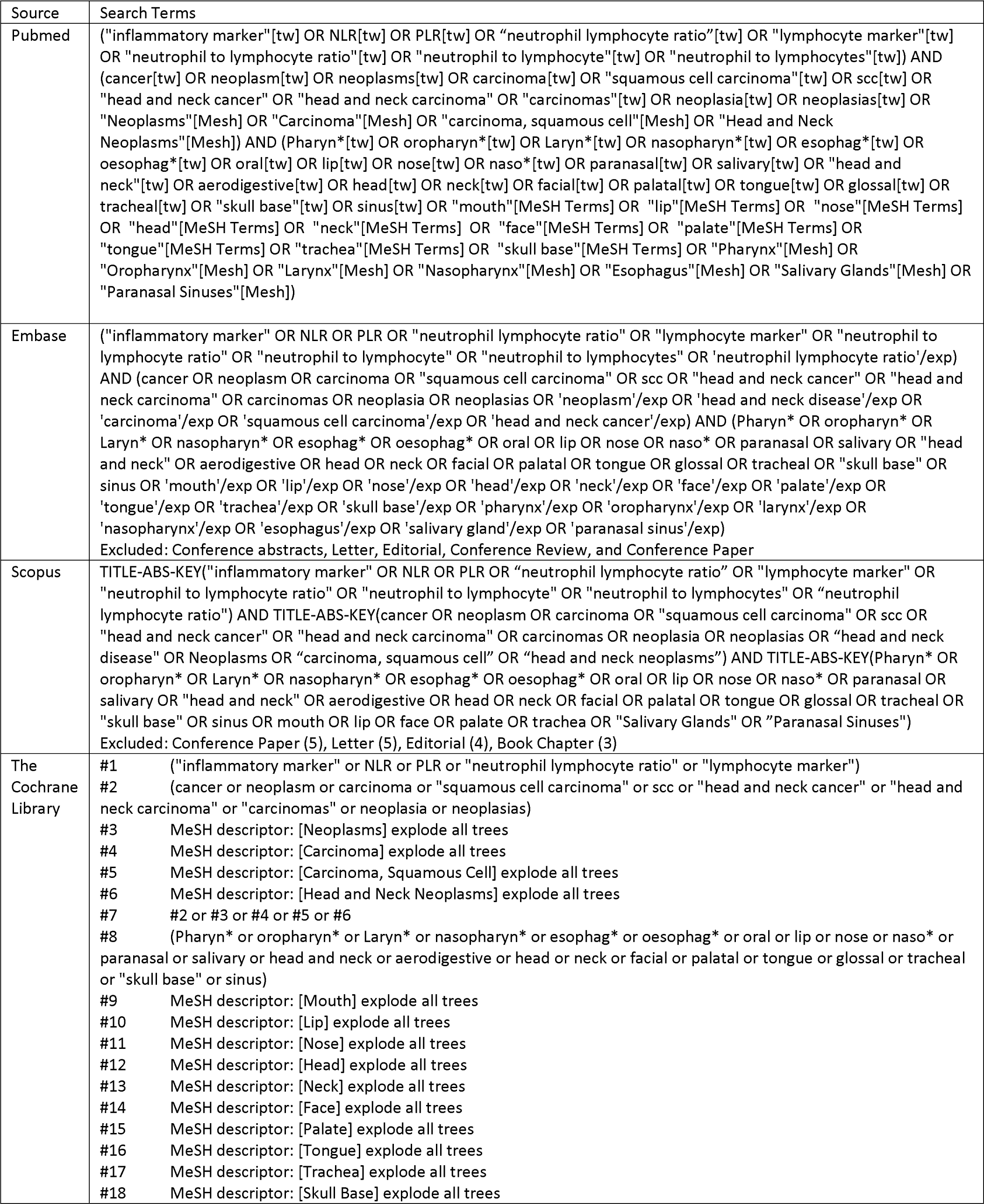
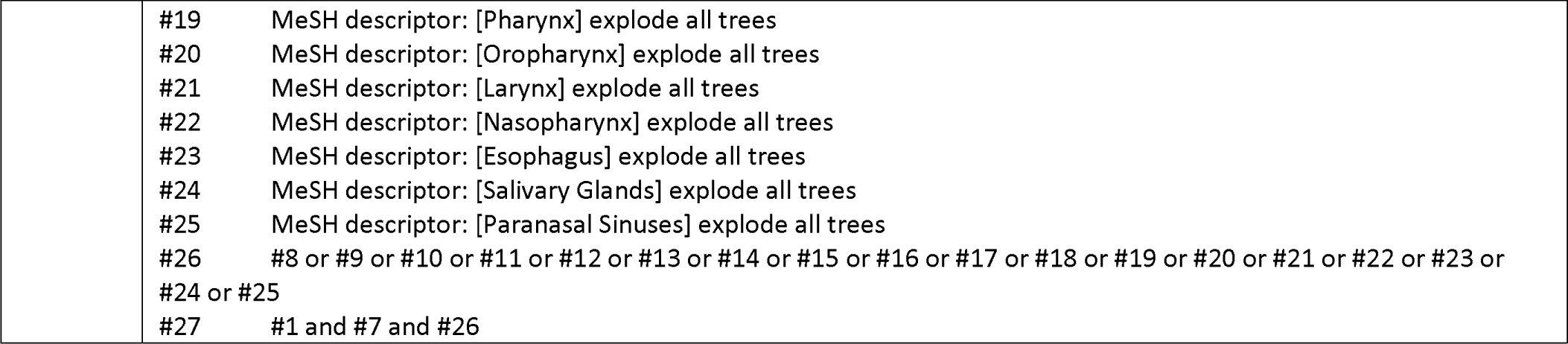
Search Strategies

### Article Selection

Articles were selected independently by two of the authors (TT, YB) in two phases. In the first phase we screened a list of titles and abstracts for full-text retrieval. During the first phase (title and abstract screening), our inclusion criteria was any study that reported a description of NLR in head and neck cancer, either in the title or abstract. If the content of the abstract was not clear, we selected the study for full-text review. Articles that passed the first phase of screening were selected for full-text retrieval, and were assessed in a second phase of screening.

In the second phase we screened full text articles using pre-determined inclusion/exclusion criteria. ^14^ Disagreements were resolved via consensus. For the second phase of the screening (full text retrieval), the following inclusion and exclusion criteria were applied. Inclusion criteria: (1) Article reports on prognostic impact of peripheral blood NLR in head and neck cancer and associated subsites; (2) NLR treated as categorical variable; (3) NLR collected prior to treatment; (4) NLR Hazard Ratio (HR) / Risk Ratio (RR) for Overall Survival (OS), with or without Disease Free Survival (DFS), with or without Progression free survival (PFS); (5) 95% Confidence interval (CI) for survival statistic, with or without the p-value; (6) Available as full text publication; (7) English Language; (8) Clinical trial, cohort, case control. Exclusion criteria: (1) Case report, conference proceeding, letters, reviews/meta analyses; (2) Thyroid and endocrine tumors; (3) Animal studies; (4) Laboratory studies; (5) Duplicate literature and duplicate data; when multiple reports describing the same population were published, only the most recent or complete report was included; (6) Metastatic cancers only; (7) Incomplete data (No NLR HR for OS). Studies with incomplete data (for example, studies that included Kaplan-Meier curves only, or without HR with 95% CI), were not excluded initially. In these cases, we contacted the corresponding authors in attempt to obtain their original data ^15–17^.

### Quality assessment

Two authors (TT, YB) jointly assessed the risk of bias in the included papers. Previous meta-analyses of observational studies have frequently used the Newcastle-Ottawa Scale (NOS) to score the risk of bias ^18^, however the NOS was not designed and validated for observational prognostic studies. Our requirement for the tool used in our paper was that it needed to be validated, widely accepted, and designed specifically for observational prognosis studies. Therefore the assessment was made using the Quality In Prognosis Studies Tool (QUIPS). ^19^ QUIPS is based on six domains: study participation, attrition, prognostic factor measurement, outcome measurement, study confounding, and statistical analysis and presentation. Each domain contained a checklist of three to nine subdomains, which were used to render a score of low, moderate or high risk of bias for the entire main domain. A detailed breakdown of the scoring criteria and subdomains may be found in the supplementary materials. Before performing the quality assessment, a training session was conducted to ensure both investigators would apply the same standards using QUIPS. Disagreements in scoring the domains were easily reconciled and consensus was reached in all assessments.

### Data Extraction

Data forms were developed *a priori* as recorded in the PROSPERO registry.^14^ Two authors (TT, YB) jointly reviewed all of the full text articles together for the data extraction process. If there were disagreements about data points, a third author (PC) was consulted to adjudicate and resolve the disagreement. The following data points were collected: First author’s name; Year of publication; Country (region) of the population studied; Sample size; Age; Gender; Demographic data; Follow-up period; Tumor data including histology, stage, grade and metastasis; Survival data HR/RR OS, RFS, DFS, PFS, with the associated 95% CI, p-value; Survival data reported with univariate or multivariate analysis; Cut-off value used to define “elevated NLR”; Method of obtaining the cut-off value; Subgroup and covariate information.

For the analysis of the relationship between NLR and clinicopathological parameters, HR/RR and 95% CI were combined as the effective value. If several estimates of NLR HR for OS were reported in the same article, we chose the most powerful one (multivariate analysis was superior to univariate analysis, and the latter one weighted over unadjusted Kaplan–Meier analysis). If the method of NLR cutoff was by done by dividing the continuous NLR data into percentile cutoffs, the highest NLR percentile cutoff was chosen for data extraction. We attempted contacting authors if the information in their paper was not sufficiently detailed to be extracted: such as details on adjusted regression analysis ^20^, or information on NLR cutoff or method of obtaining the cutoff. ^20–22^ If the HR for OS was reported as HR of a patient with NLR *below* a specific cutoff experiencing the endpoint of death (versus HR of a patient with NLR *above* a specific cutoff experiencing endpoint of death), we took the reciprocal of the reported HR in order to make it comparable to the other studies. ^22–25^

## Statistical Analysis

The logarithm of the HR with Standard Error (SE) was used as the primary summary statistic. To obtain the log[HR] and SE, the HR with 95% CI was extracted directly from the studies. Additional calculation to obtain the HR was required if the study reported the reciprocal of the HR. Estimates of log[HR] were weighted and pooled using the generic inverse-variance.^11^ Because of anticipated heterogeneity, a more conservative approach applying the random effects model (DerSimonian and Laird method) was chosen for all analyses. Forest plots were constructed for all outcomes displaying the random-effects model of the summary effect measure and 95% CI. Heterogeneity was assessed using Cochran’s Q and Higgins’s I^2^. Cochrane’s Q p-value of <0.1 and I^2^ > 50% were considered as markers of significant heterogeneity. To assess publication bias, Begg’s Funnel Plot and Egger’s bias indicator test were used. If publication bias was detected, the influence of bias on the overall effect was assessed by Duval’s “Trim and fill” method. ^26^ A Failsafe N measure was also calculated with the methods described by Rosenthal.^27^ All analyses was done using the RevMan 5.3 analysis software (Cochrane Collaboration, Copenhagen, Denmark).^28^ Tests for publication bias were performed by Meta-Essentials (ERASMUS Research Institute, Rotterdam, Netherlands). ^29^ All statistical tests were two-sided, and a p-value of less than 0.05 was considered statistically significant. No correction was made for multiple testing.

## Results

### Study Characteristics

The PRISMA flow chart of the systematic review can be found in Figure 1. An initial search done using the search strategy (Table 1) obtained an initial 900 results.

**Figure 1.**
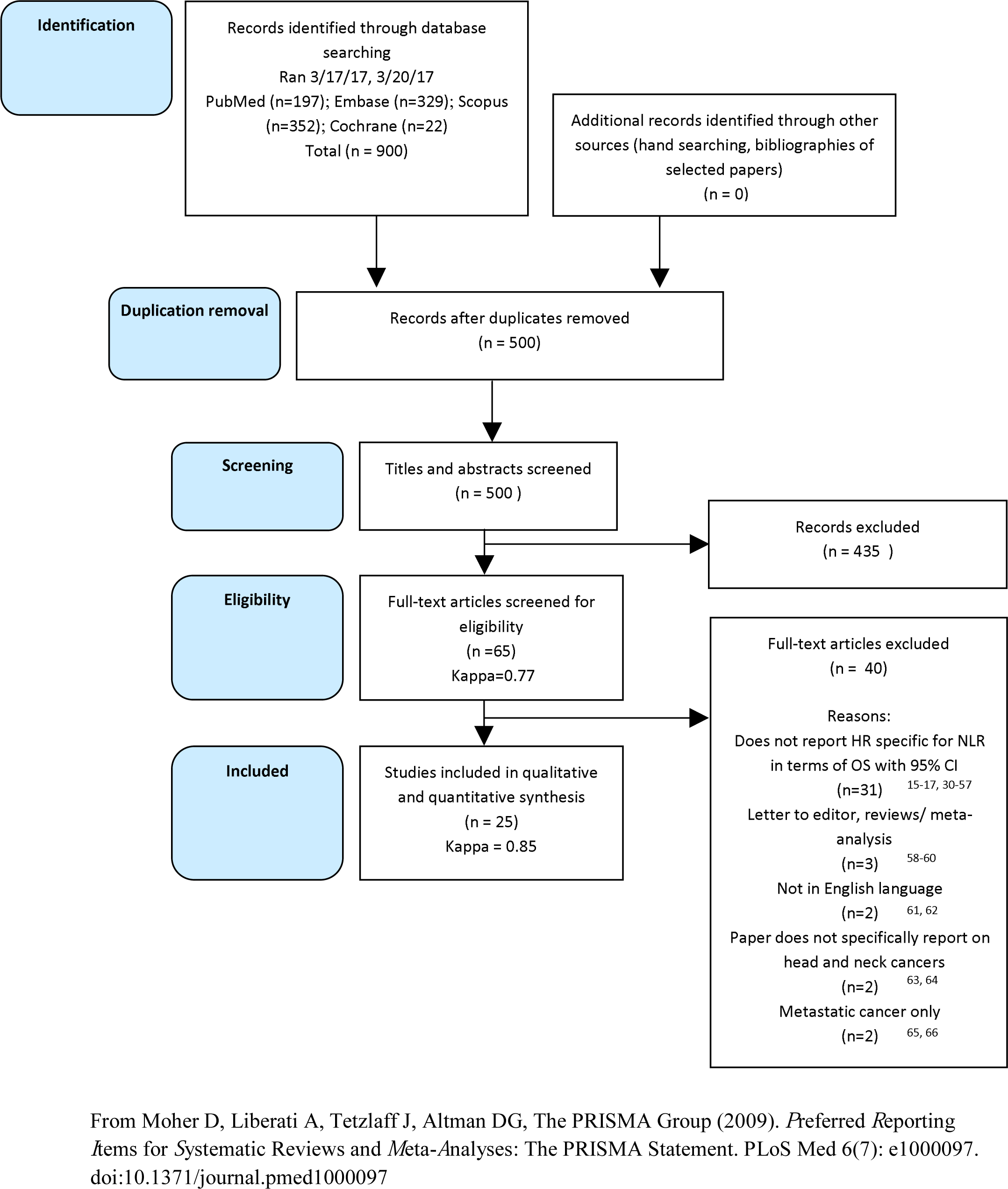
PRISMA Flowchart

De-duplication was then performed, which reduced the number of results to 500. The first phase of screening was performed next on titles and abstracts, which reduced the number of results to 65. The agreement was good for the first phase with a Kappa of 0.7. The second phase of screening resulted in the exclusion of a further 40 results. The agreement was very good for the second phase of screening with a Kappa of 0.85. The list of excluded papers with the reasons for exclusion may be found in the Supplementary Materials.

Thus, 25 studies published between 2011 and 2017 were included in our meta-analysis, with sample sizes ranging from 59 to 1410 patients. ^20–25, 36, 67–84^As the study by Charles et al^68^ had split their data into two cohorts with separately reported HR and 95% CI, we have designated both groups of data as Charles1 and Charles2 respectively. The characteristics of the included studies are summarized in Table 2. Eleven studies were from China, three from Japan, two from Korea, two from the USA, one from Australia (two cohorts), one from Austria, one from India, one from Italy, one from Singapore and one from the UK. Out of 25 studies, one was a prospective cohort study. The rest of the studies were based on retrospectively collected data.

**Table 2.**
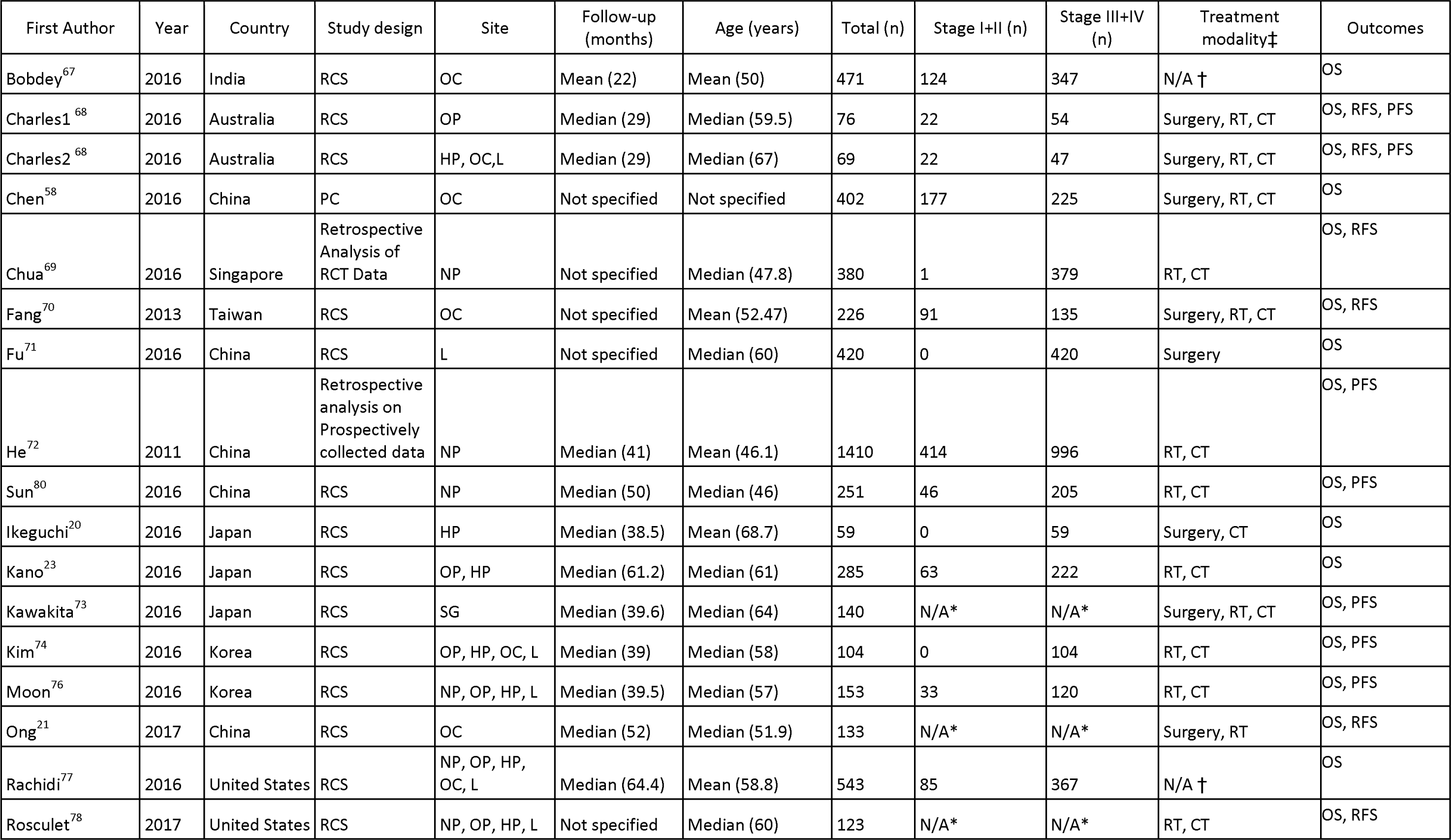
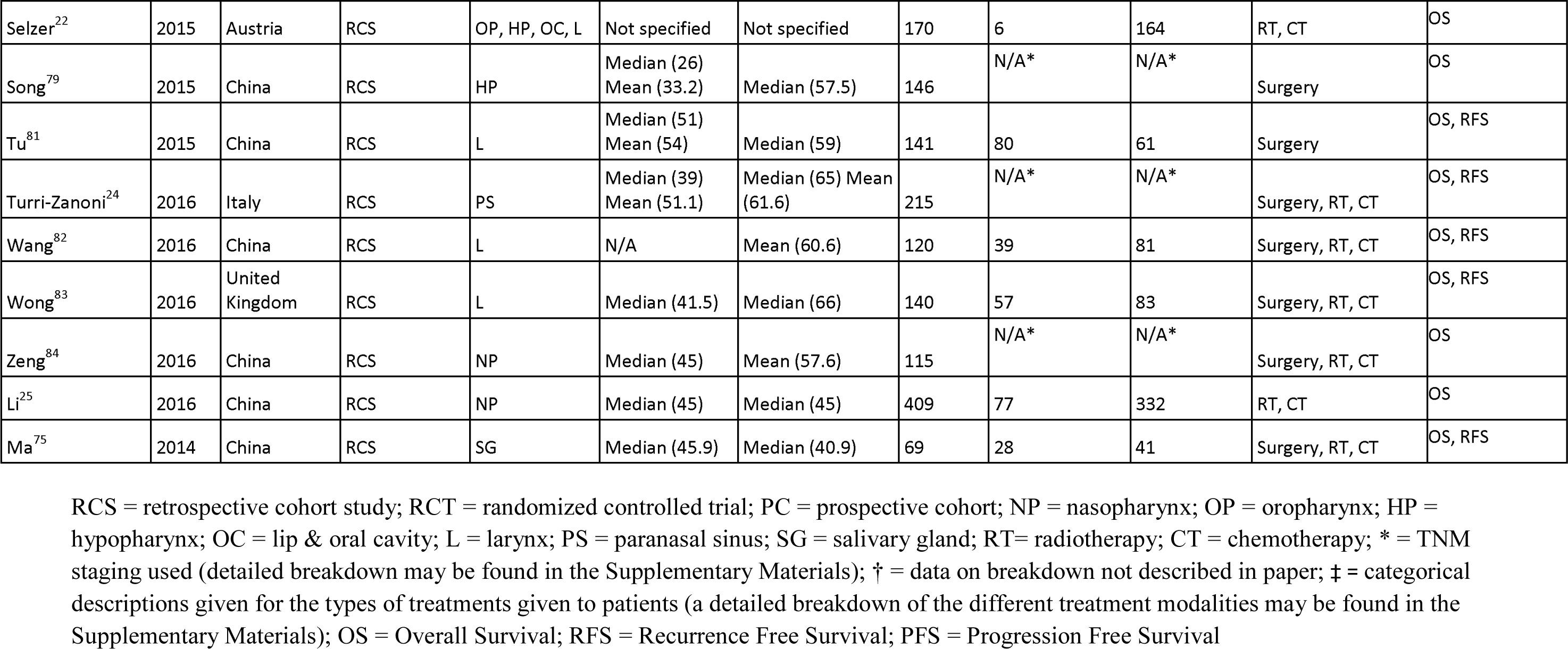
Summary of Included Studies

NLR was calculated from laboratory data in all of the studies. NLR cutoffs ranged from 1.92 – 5.56 (Median 2.895), with cutoffs unavailable from two studies. (Table 3) NLR cutoffs were obtained from ROC curve analysis or training sets in 11 studies, based on previous literature in 4 studies, median value in 4 studies, and percentile in 3 studies, and not mentioned in 3 studies. (Table 3) HR and 95% CI was reported in the original literature in 23 of the studies, and converted from the reciprocal in 2 studies. HR in 24 out of 25 studies was available through multivariate analysis (Table 3). The full data extracted from the papers can be found in the Supplementary Materials.

**Table 3.**
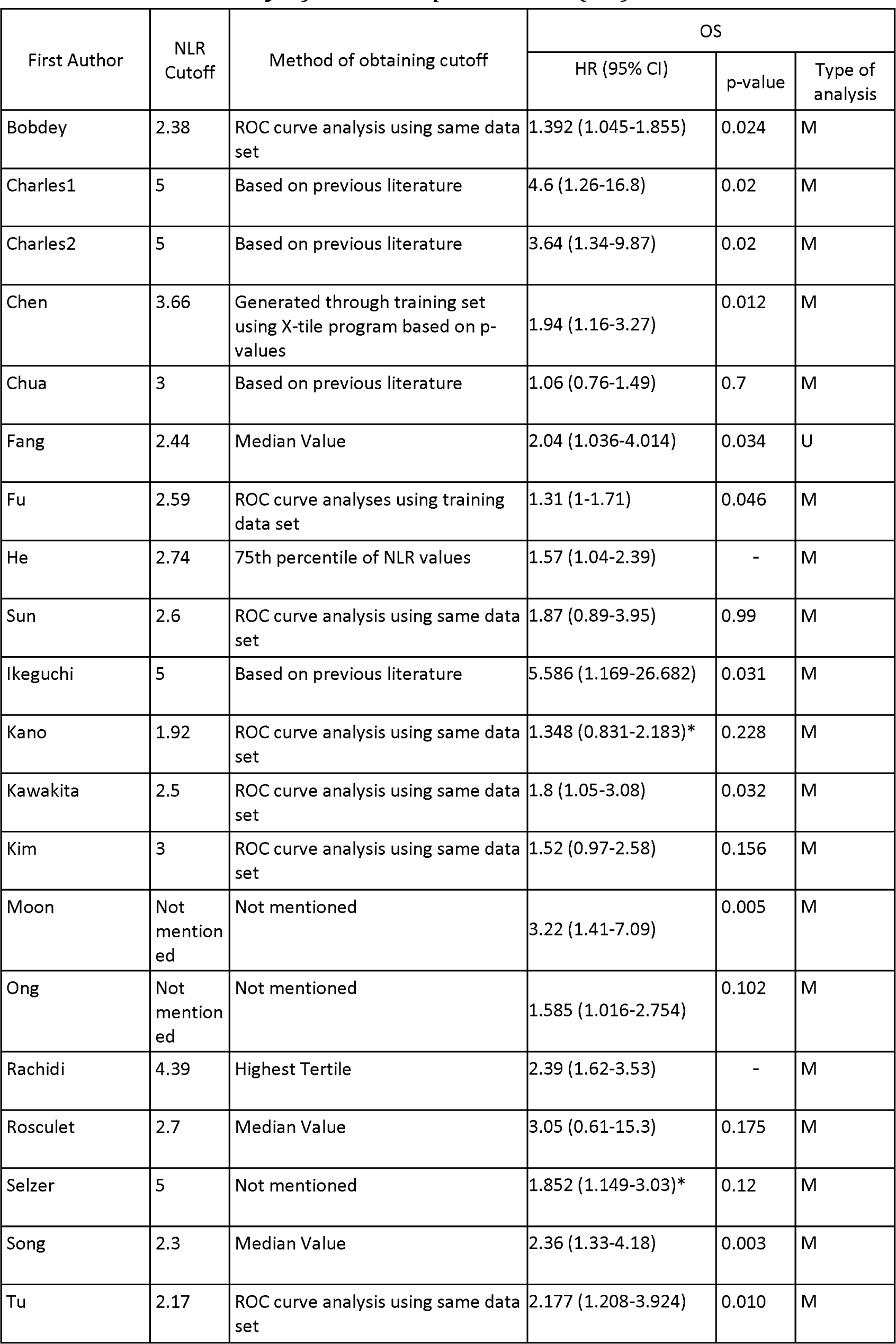
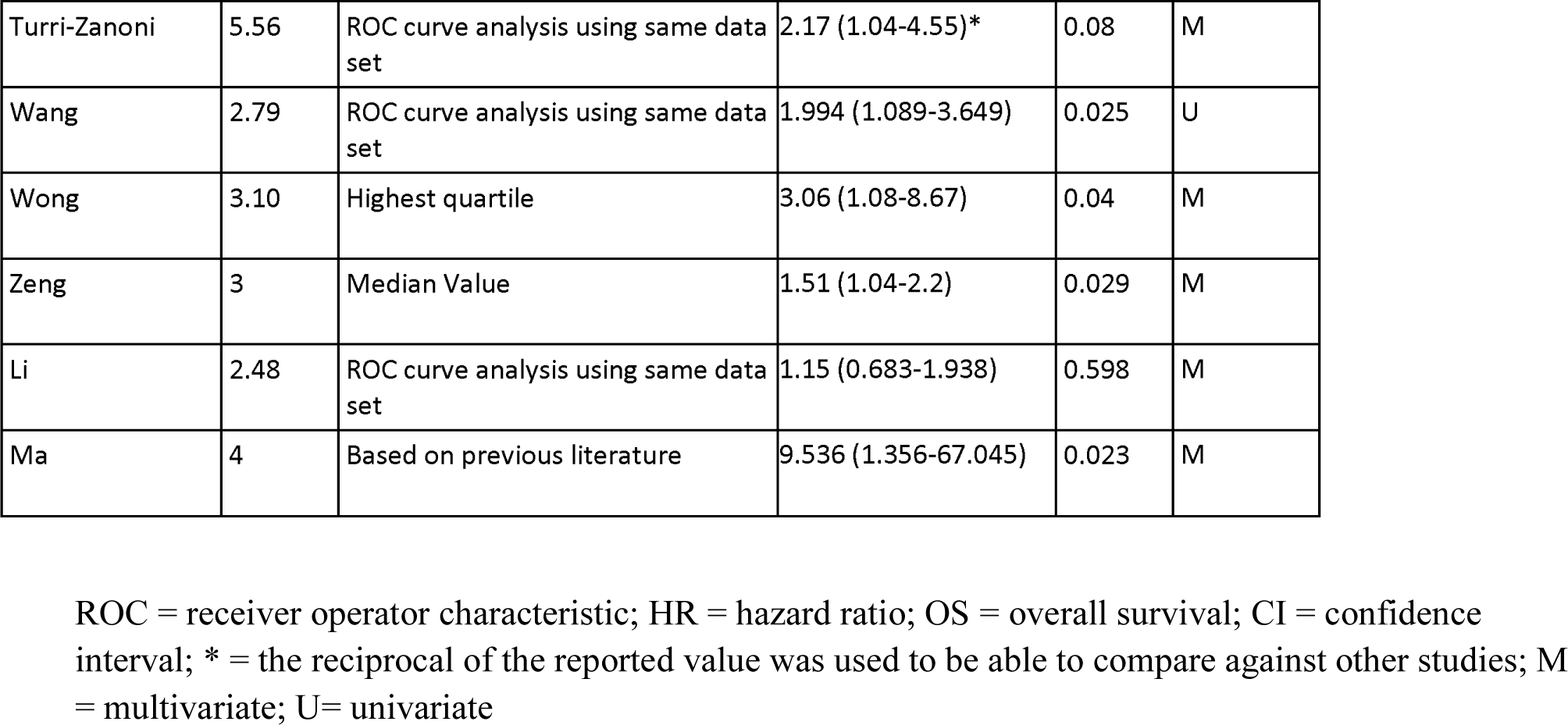
Summary of NLR Endpoint Data (OS)

### Quality Assessment

The included studies were at low to moderate risk of bias with regards to study participation and study attrition. As a large majority of the studies were retrospective cohort studies, we found that there was an inherent risk of bias in patient selection. There were also studies that did not adequately report the attrition rate, or numbers of patients who were excluded because of unavailable data. Prognostic factor measurement (method of obtaining the NLR cutoff) for most of the studies was at a moderate risk of bias. A number of the selected studies had derived their NLR cutoff values from previous literature, from percentile values of NLR, or was not even reported. There were some high quality studies that derived the NLR cutoff from Receiver Operator Characteristic (ROC) curve analysis alone or with a ‘training cohort’ data set. Outcome measurement was mostly at a low risk of bias for the selected studies, with many studies having clear definitions and descriptions of their endpoints. Study confounding was at a low to moderate risk of bias, with most studies having appropriate and sufficient covariates. Statistical analysis and reporting was mostly at a low risk of bias, with most studies having the appropriate statistical designs and data reporting. There was only a minority of studies that only reported univariate/unadjusted data.

Figure 2 shows the overall summary of the quality assessment grading, and Table 4 shows the grading at the level of the individual study. (See Supplementary files for detailed QUIPS criteria)

**Table 4.** Study-Level Quality Assessment Using the Quality In Prognosis Studies Tool (QUIPS)

| Study | Study participation | Study attrition | Prognostic factor measurement | Outcome measurement | Study confounding | Statistical analysis and reporting |
| --- | --- | --- | --- | --- | --- | --- |
| Bobdey | H | H | H | L | M | M |
| Charles | M | L | M | L | L | L |
| Chen | L | L | L | L | L | L |
| Chua | L | M | M | L | L | L |
| Fang | M | M | M | L | M | L |
| Fu | L | M | L | L | L | L |
| He | M | M | M | L | M | M |
| Sun | M | L | M | L | M | L |
| Ikeguchi | M | M | M | M | M | H |
| Kano | M | M | M | M | M | L |
| Kawakita | H | H | M | L | L | L |
| Kim | L | L | M | L | L | H |
| Moon | L | L | H | M | L | H |
| Ong | M | M | H | M | L | H |
| Rachidi | L | M | M | M | L | L |
| Rosculet | L | L | M | L | L | L |
| Selzer | L | M | H | M | L | M |
| Song | M | M | M | M | L | L |
| Tu | M | M | M | L | L | L |
| Turri-Zanoni | L | L | M | L | L | L |
| Wang | M | L | M | L | M | M |
| Wong | L | M | M | M | M | L |
| Zeng | M | L | M | L | L | L |
| Li | M | H | M | L | M | L |
| Ma | L | L | M | L | L | L |

H, high risk of bias; L, low risk of bias; M, moderate risk of bias.

**Table 5.**
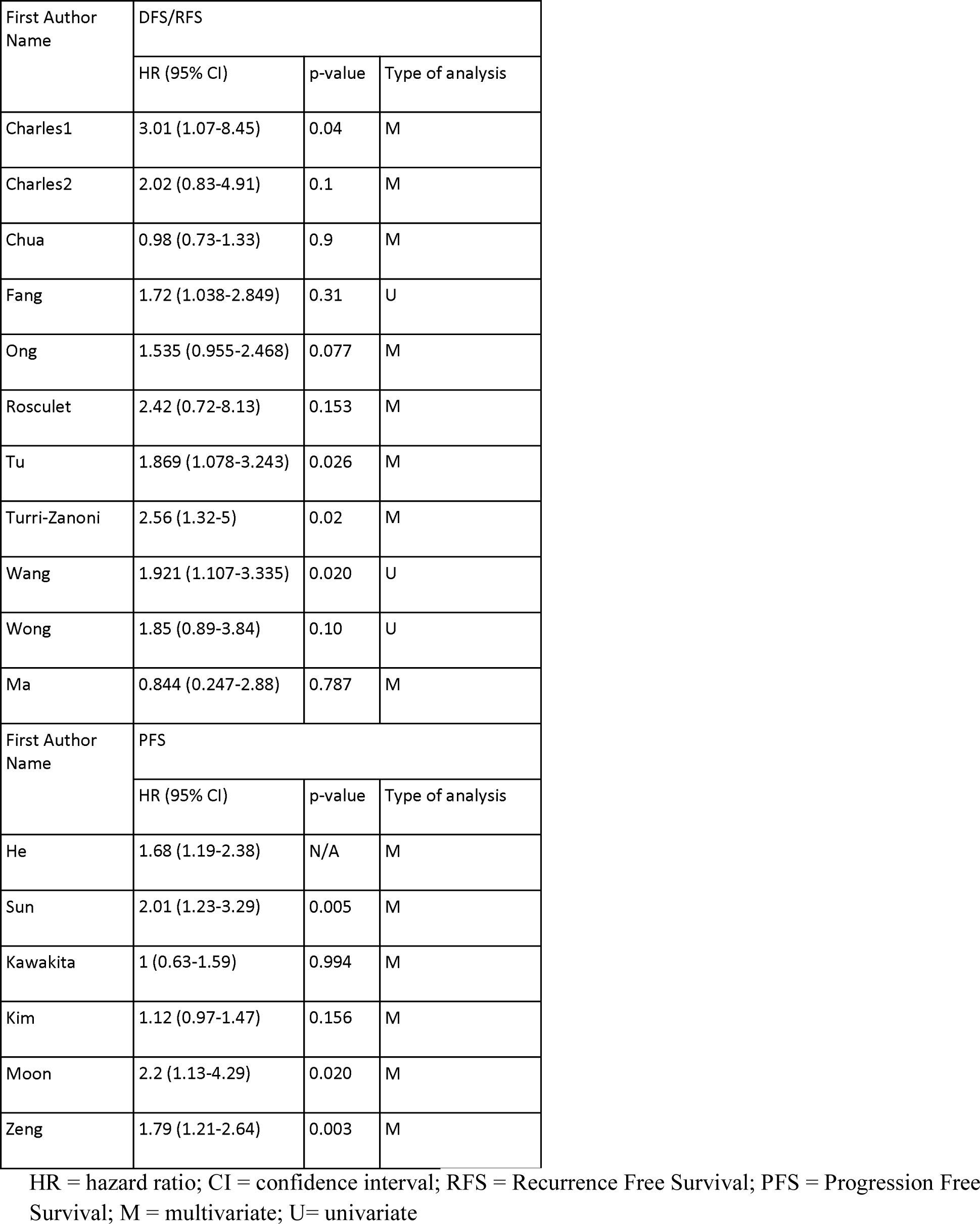
DFS/RFS and PFS from selected studies

**Figure 2.**
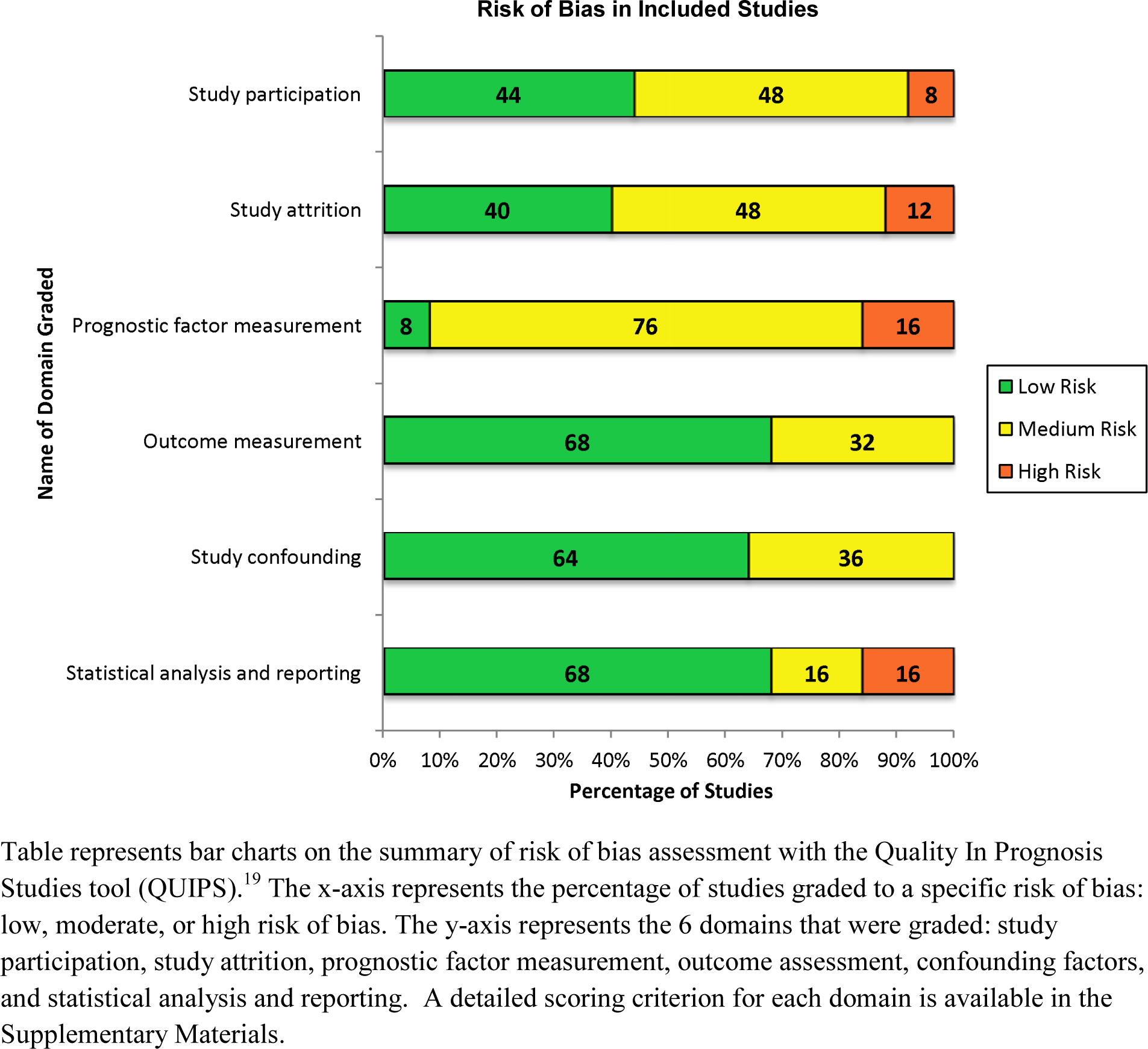
Risk of Bias Summary

### NLR and OS in HNC

Data from 25 studies (26 cohorts) were synthesized in the meta-analysis for NLR and OS in HNC patients. An elevated NLR value was found to be significantly associated with poorer OS with HR of 1.85 (95% CI: 1.58-2.17, p<0.00001). The test for heterogeneity showed an I^2^ value of 59%, and p-value<0.0001, which represented significant heterogeneity of the results. The Forest Plot and corresponding Funnel Plot are represented in Figure 3.

### NLR and DFS/RFS/PFS in HNC

Data from 11 studies (12 cohorts) were synthesized in the meta-analysis for NLR and DFS/RFS in HNC. An elevated NLR value above the cutoff was found to be significantly associated with poorer DFS/RFS with HR of 1.66 (95% CI: 1.30-2.11, p<0.0001). The test for heterogeneity showed an I^2^ value of 38%, and p-value of 0.1, which represented marginal heterogeneity of the results. (Figure 4)

Data from 5 studies were synthesized in the meta-analysis for NLR and PFS in HNC. An elevated NLR value above the cutoff was found to be significantly associated with poorer PFS with HR of 1.43 (95% CI: 1.09-1.87, p=0.001). The test for heterogeneity showed an I^2^ value of 69%, and p-value of 0.01, which represented substantial heterogeneity of the results. (Figure 5)

### Subgroup Analysis for NLR and OS

To explore the heterogeneity of NLR and OS, subgroup analysis was performed for tumor site, treatment type, geographical location, sample size, method of obtaining NLR cutoff and NLR cutoff value. Subgroup analysis for tumor site and treatment type was decided *a priori*, as listed in our online protocol^14^. Subgroup analysis for the rest of the parameters was decided *post hoc*. Unfortunately because of limited data, subgroup analysis according to TNM staging or tumor grade was not performed. Due to the small number of studies representing DFS/RFS and PFS, subgroup analysis of these measures was not performed to avoid statistical instability.

### NLR and Tumor Site in HNC

Tumor sites were grouped according to their shared biology and etiology: ^85^ Oral cavity (OC), oropharyngeal (OP), and hypopharyngeal (HP) cancers were grouped together; Nasopharyngeal (NP) cancers and Laryngeal (L) cancers were grouped separately; Salivary cancers and others were grouped together as “Other”. Because of the unavailability of individual patient information in many of the studies, cohorts that had reported sites of tumors across multiple of the aforementioned categories were not included in this subgroup analysis. Data from 19 studies were included in this subgroup analysis. The HR for each group indicated significant association between elevated NLR and OS: For the OC/OP/HP group the HR was 1.83 (95% CI: 1.43-2.35, p<0.00001); for the NP group the HR was 1.28 (95% CI: 1.01-1.62, p=0.04); for the L group the HR was 1.60 (95% CI: 1.27-2.03, p<0.0001); for the Other group the HR was 2.17 (95% CI: 1.27-3.71, p=0.004). Heterogeneity was low within all subgroups with I^2^ values in the OC/OP/HP, NP, L, and Other groups of 27%, 10%, 21% and 24% respectively. However, test for differences between subgroups showed homogeneity (I^2^=48.3%, p=0.12). (Table 6)

**Table 6.** Subgroup Analysis

| Stratified analysis | Number of Cohorts* | Random Effects Model |  | Heterogeneity |  |
| --- | --- | --- | --- | --- | --- |
|  |  | HR (95% CI) | p | I<sup>2</sup>(%) | Ph |
| Subgroup 1: Tumor Site |  |  |  |  |  |
| OC, OP, HP | 7 | 1.83 (1.43-2.35) | <0.00001 | 27 | 0.22 |
| NP | 4 | 1.28(1.01-1.62) | 0.04 | 10 | 0.34 |
| L | 5 | 1.60(1.27-2.03) | <0.00001 | 21 | 0.28 |
| Other | 3 | 2.17(1.27-3.71) | 0.004 | 24 | 0.27 |
|  |  |  |  | Test for subgroup differences |  |
|  |  |  |  | 48 | 0.12 |
| Subgroup 2: Treatment Type |  |  |  |  |  |
| Surgical | 3 | 1.76(1.16-2.68) | 0.008 | 59 | 0.09 |
| Non-surgical | 9 | 1.48(1.21-1.80) | 0.0001 | 22 | 0.25 |
|  |  |  |  | Test for subgroup differences |  |
|  |  |  |  | 0 | 0.45 |
| Subgroup 3: Geographical Location |  |  |  |  |  |
| Asian Ethnicity | 19 | 1.57(1.38-1.79) | <0.00001 | 23 | 0.18 |
| Caucasian Ethnicity | 7 | 2.72(2.13-3.47) | <0.00001 | 21 | 0.27 |
|  |  |  |  | Test for subgroup differences |  |
|  |  |  |  | 93.4 | <0.00001 |
| Subgroup 4: Sample Size |  |  |  |  |  |
| Small Sample | 13 | 2.22(1.72-2.87) | <0.00001 | 54 | 0.01 |
| Large Sample | 13 | 1.58(1.34-1.85) | <0.00001 | 38 | 0.08 |
|  |  |  |  | Test for subgroup differences |  |
|  |  |  |  | 80 | 0.03 |
| Subgroup 5: Method of NLR Cutoff |  |  |  |  |  |
| ROC Curve/Training Set | 11 | 1.51(1.32-1.73) | <0.00001 | 0 | 0.68 |
| Other Method | 12 | 2.26(1.63-3.13) | <0.00001 | 73 | <0.0001 |
|  |  |  |  | Test for subgroup differences |  |
|  |  |  |  | 80 | 0.03 |
| Subgroup 6: NLR Cutoff |  |  |  |  |  |
| NLR Cutoff<2.895 | 12 | 1.53(1.34-1.75) | <0.00001 | 0 | 0.58 |
| NLR Cutoff>2.895 | 12 | 2.15(1.58-2.91) | <0.00001 | 73 | <0.0001 |
|  |  |  |  | Test for subgroup differences |  |
|  |  |  |  | 74 | 0.05 |

### NLR and Treatment Type

Treatment type was stratified into surgical and non-surgical. Because of the unavailability of individual patient information in some of the studies, cohorts that had reported surgical plus non-surgical therapies were not included in this subgroup analysis. Data from 12 studies were included in this subgroup analysis. The HR for each group indicated significant association between elevated NLR and OS: For the surgery-only group the HR was 1.76 (95% CI: 1.16-2.68, p=0.008); for the non-surgical group the HR was 1.48 (95% CI: 1.21-1.80, p=0.0001). Heterogeneity was high in the surgery group and low in the non-surgery group with I^2^ values of 59% and 22% respectively. The test for differences between subgroups showed homogeneity (I^2^=0%, p=0.45). (Table 6)

### NLR and Geographical Location (Ethnicity)

Geographical location was stratified according to predominantly Asian populations versus predominantly Caucasian populations. The HR for each group indicated significant association between elevated NLR and OS: For the Asian ethnicity group the HR was 1.57 (95% CI: 1.38-1.79, p<0.00001); for the Caucasian ethnicity group the HR was 2.72 (95% CI: 2.13-3.47, p<0.00001). Heterogeneity was low within the Asian and Caucasian groups with I^2^ of 23% and 21% respectively. The test for differences between subgroups was significant (I^2^=93%, p<0.0001). (Table 6)

### NLR and Study Size

The size of a study was stratified according to the median value of the sample size in the studies selected for meta-analyses (Median sample size was n=146 patients). Small studies had n ≤ 146 patients, and large samples had n > 146 patients. The HR for each group indicated significant association between elevated NLR and OS: For the small studies, the HR was 2.22 (95% CI: 1.72-2.87, p<0.00001); for the large studies, the HR was 1.58 (95% CI: 1.34-1.85, p<0.00001). Heterogeneity within the small and large studies was moderate-significant, with I^2^ values of 54% and 38% respectively. The test for differences between subgroups was significant (I^2^=80%, p=0.03). (Table 6)

### NLR and Method of NLR Cutoff

Method of obtaining NLR cutoff was stratified into ROC curve analysis or training sets, versus other methods of obtaining NLR cutoff. Data from 23 cohorts were included in this analysis. The HR for each group indicated significant association between elevated NLR and OS: for the ROC curve/training set group the HR was 1.51 (95% CI: 1.32-1.73, p<0.00001); for the Other Methods group the HR was 2.26 (95% CI: 1.63-3.13, p<0.00001). Heterogeneity was low in the ROC/training set group but high in the Other Methods group, with I^2^ of 0% and 73%, respectively. The test for differences between subgroups was significant (I^2^=79.8%, p=0.03). (Table 6)

### NLR and Cutoff Value

Subgroup analysis for NLR cutoff values was performed by comparing studies above and below the median NLR cutoff value (median NLR cutoff was 2.895). Data from 24 cohorts were included in this analysis. The HR for each group indicated significant association between elevated NLR and OS: for the NLR cutoff<2.895 group the HR was 1.53 (95% CI: 1.34-1.75, p<0.00001); for the NLR cutoff>2.895 group the HR was 2.15 (95% CI: 1.58-2.91, p<0.00001). Heterogeneity was low in the NLR<2.895 group and high in the NLR>2.895 group with I^2^ values of 0% and 73% respectively. The test for differences between subgroups significant (I^2^=74.3%, p=0.05). (Table 6)

### Sensitivity Analysis

A single study involved in the meta-analysis was deleted each time to unveil the influence of the individual data set to the pooled HRs for OS, and the corresponding pooled HR for OS was not significantly changed (data not shown).

### Publication Bias

Begg’s Funnel Plot for HR of OS indicated that there was evidence of publication bias, with fewer negative small studies reporting negative results than would be expected. (Figure 3) The p value for Egger’s test indicated there was publication bias for OS (p=0.001) but not for DFS/RFS (p=0.17) or PFS (p=0.45). Therefore, we further performed Duval’s “trim and fill” analysis for OS data. It was estimated that an additional 9 studies evaluating the prognostic value of NLR remain unpublished. The filled meta-analysis for the effect of NLR in OS upheld our pooled results (adjusted HR: 1.55, 95% CI: 1.30-1.85, p<0.00001). A classic Failsafe N value was also calculated which showed that an additional 587 negative studies are needed to invalidate the results of the meta-analysis of OS data. (Supplementary Materials)

## Discussion

This meta-analysis aimed to examine the relationship between NLR and OS, DFS/RFS, and PFS in HNC. Our analysis for OS combined the results of over 6770 patients in 26 cohorts (25 studies). The pooled data demonstrated that an elevated pretreatment NLR significantly predicted poorer OS (HR: 1.85, 95% CI: 1.58-2.17), DFS/RFS (HR: 1.43, 95% CI: 1.30-2.11), and PFS (HR 1.43, 95% CI: 1.09-1.87) of HNC patients. There was heterogeneity of results for all of the above endpoints. Although there was significant heterogeneity of results for OS (I^2^=59%), the prognostic significance was not weakened by subgroup analysis stratified by tumor site, treatment type, geographical location (ethnicity), sample size, method of obtaining NLR cutoff and NLR cutoff value. The within-subgroup heterogeneity for OS was eliminated in subgroup analysis of individual tumor sites, in non-surgical treatments, within specific ethnicities, within studies using ROC curve/training set analyses, and those with NLR cutoff <2.895. (Table 6) Of note, when comparing between subgroups for differences, statistically significant differences were found for ethnicity (I^2^=93%, p<0.00001), study size (I^2^=80%, p=0.03), method of NLR cutoff (I^2^=0%, p<0.00001), and NLR cutoff of 2.895 (I^2^=74%, p=0.05) (Table 6). In these subgroups, both Cochran’s Q and Higgins’s I^2^ indicated a significant interaction existed between the subtotal estimates for the subgroups. Thus, it can be concluded that these subgroup stratifications estimated different population parameters.

The low within-group heterogeneity in the tumor site and ethnicity subgroups may point to different NLR cutoffs existing for each patient population. This is clinically consistent as we would expect tumors of different subsites and from different patient populations to have different characteristics that would affect survival. The high between-group heterogeneity and low within-group heterogeneity in subgroups analyzing study design (sample size, NLR cutoff method, NLR cutoff) could also point to the conclusion that higher quality studies had different results from lower quality studies. These differences were shown to be quantitative in nature, as the differences of effect were still in the same direction. It was expected that studies using more robust statistical techniques would come to a similar conclusion, as demonstrated by the low within group heterogeneity of studies using ROC curve analysis (I^2^=0%). These findings suggest that the effect of a dichotomized cutoff for NLR may have utility in different populations, and could be used to guide clinical stratification and decision making with regard to outcomes for HNC patients. Notwithstanding, due to the intrinsic limitations to meta-analyses, we recommend prudence to avoid over interpreting the results of the subgroup analysis. To the best of our knowledge, this is the first meta-analysis reporting the relationship between elevated pretreatment NLR and outcomes in all sites of the head and neck.

The paradigm of local and systemic inflammatory states interacting with the local tumor microenvironment is based on strong evidence. ^2, 86^ However, the mechanism behind the association between a high NLR and poor cancer prognosis remain poorly understood. ^5^A high NLR indicates a relative neutrophilia and lymphopenia, and neutrophilia has been known to inhibit the cytolytic activity of T cells and NK cells.^87, 88^ On the other hand, the significance of lymphocyte infiltration of tumors has been shown to improve prognosis and response to treatment.^89^ Perhaps the prognostic ability of NLR lies in its measurement of the pro-tumor versus anti-tumor dynamic in the host immune system.^5^ The prognostic utility of NLR has been demonstrated in other cancers, including hepatocellular carcinoma^7^, lung cancer^90^, urinary tract cancer^6^, colorectal cancer^91^, renal^92^, and others^5^. Our study results are consistent with results from these previously reported meta-analyses in other cancers. A search on the clinicaltrials.gov database has also shown that there is a prospective clinical trial investigating NLR in HNC that is already underway (NCT02211677). Recently, novel staging systems and prognostic scoring systems have also been developed for HNC incorporating NLR, such as using platelets and NLR (COP-NLR) ^31^, or histopathological staging and NLR^42^. Other inflammatory markers and systems such as the platelet to lymphocyte ratio (PLR) ^24^, lymphocyte to monocyte ratio (LMR) ^23^and Glasgow Prognostic Score (GPS)^93^ have also received interest as prognostic indicators in HNC. It remains to be seen which of these markers, or combination of markers, is the superior option for clinical use as a prognostic biomarker.

In spite of our findings, there are several weaknesses of our study that we acknowledge. Heterogeneity was found in the pooled results for OS, PFS, and marginally so for DFS. Subgroup analysis on OS showed that tumor site, ethnicity and study design factors could account for the heterogeneity. It is very likely that the heterogeneity is secondary to the above factors, together with the unreported genetic diversity of head and neck cancers as well as other confounders (such as HPV status). Most of the studies included were also retrospective in nature, with only one study collecting data prospectively. Furthermore, because of a lack of individual patient data in many of the studies, we were unable to perform meta-analyses of individual patient data (MAIPD). We were also unable to include all cohorts in the subgroup analysis due to the diverse patient populations represented in the included studies. Another limitation of this paper is the publication bias detected for OS, as there were significantly more papers published that reported a poorer OS for higher NLR. However, the adjusted trim and fill analysis did not change the original conclusion. Lastly, the primary endpoint chosen for inclusion of studies was OS, therefore DFS and PFS data were drawn from studies that reported OS as an endpoint.

The advantages of our study were the relatively high amount of studies included, agreement of our results with the existing literature in other cancers, and significance using the random effects model. The effect of NLR on OS was also stable after performing subgroup analysis, sensitivity analysis, and “trim and fill” publication bias adjustment. The quality of studies that were included also showed low-moderate bias as assessed via QUIPS, with only few studies having a high amount of bias.

To conclude, the results of our meta-analysis suggest that an elevated pretreatment NLR is a negative prognostic factor in patients with HNC. The NLR value could have utility in stratifying patients and determining patient-specific treatment plans, particularly identifying high-risk patients that might benefit from adjuvant therapy. Our results should be interpreted with some degree of caution in view of the limitations described above. Therefore, further research with high-quality prospective studies is needed to fully validate the prognostic utility of NLR in HNC.

## Acknowledgement

We would like to sincerely thank Janice Lester, MLS from Long-Island Jewish Medical Center Hospital for reviewing and providing advice on the final search strategy.

## References

1. Jemal A, Bray F, Center MM, Ferlay J, Ward E, Forman D. Global cancer statistics. CA: a cancer journal for clinicians. 2011;61(2):69–90.

2. Grivennikov SI, Greten FR, Karin M. Immunity, inflammation, and cancer. Cell. 2010;140(6):883–99.

3. O’callaghan DS, O’donnell D, O’connell F, O’byrne KJ. The role of inflammation in the pathogenesis of non-small cell lung cancer. Journal of Thoracic Oncology. 2010;5(12):2024–36.

4. Aggarwal BB, Vijayalekshmi R, Sung B. Targeting inflammatory pathways for prevention and therapy of cancer: short-term friend, long-term foe. Clinical Cancer Research. 2009;15(2):425–30.

5. Templeton AJ, McNamara MG, Šeruga B, Vera-Badillo FE, Aneja P, Ocaña A, et al. Prognostic role of neutrophil-to-lymphocyte ratio in solid tumors: a systematic review and meta-analysis. Journal of the National Cancer Institute. 2014;106(6):dju124.

6. Wei Y, Jiang Y-Z, Qian W-H. Prognostic role of NLR in urinary cancers: a meta-analysis. PloS one. 2014;9(3):e92079.

7. Xiao W-K, Chen D, Li S-Q, Fu S-J, Peng B-G, Liang L-J. Prognostic significance of neutrophil-lymphocyte ratio in hepatocellular carcinoma: a meta-analysis. BMC cancer. 2014;14(1):117.

8. Bakshi SS. Letter to the editor regarding neutrophil-to-lymphocyte ratio in laryngeal squamous cell carcinoma. Head & neck. 2017;39(3):614.

9. Su L, Zhang M, Zhang W, Cai C, Hong J. Pretreatment hematologic markers as prognostic factors in patients with nasopharyngeal carcinoma: A systematic review and meta-analysis. Medicine. 2017;96(11).

10. Yu B, Li Z, Zheng Q, Luo Z, Li J, Zhou Y, et al. Prognostic value of neutrophil to lymphocyte ratio in patients with nasopharyngeal carcinoma: A meta-analysis. Biomedical Research. 2017;28(3).

11. Macaskill P, Gatsonis C, Deeks J, Harbord R, Takwoingi Y. Cochrane handbook for systematic reviews of diagnostic test accuracy. Version 09 0 London: The Cochrane Collaboration. 2010.

12. Moher D, Liberati A, Tetzlaff J, Altman DG, Group P. Preferred reporting items for systematic reviews and meta-analyses: the PRISMA statement. PLoS med. 2009;6(7):e1000097.

13. Stroup DF, Berlin JA, Morton SC, Olkin I, Williamson GD, Rennie D, et al. Meta-analysis of observational studies in epidemiology: a proposal for reporting. Jama. 2000;283(15):2008–12.

14. Neutrophil-lymphocyte ratio as a predictor of prognosis in head and neck cancer: a systematic review and meta-analysis. PROSPERO 2017:CRD42017059500 [Internet]. 2017. Available from: http://www.crd.york.ac.uk/PROSPERO/display_record.asp?ID=CRD42017059500.

15. Farhan-Alanie OM, McMahon J, McMillan DC. Systemic inflammatory response and survival in patients undergoing curative resection of oral squamous cell carcinoma. The British journal of oral & maxillofacial surgery. 2015;53(2):126–31.

16. Haddad CR, Guo L, Clarke S, Guminski A, Back M, Eade T. Neutrophil-to-lymphocyte ratio in head and neck cancer. Journal of medical imaging and radiation oncology. 2015;59(4):514–9.

17. Kara M, Uysal S, Altinisik U, Cevizci S, Guclu O, Derekoy FS. The pre-treatment neutrophil-to-lymphocyte ratio, platelet-to-lymphocyte ratio, and red cell distribution width predict prognosis in patients with laryngeal carcinoma. European archives of oto-rhino-laryngology: official journal of the European Federation of Oto-Rhino-Laryngological Societies (EUFOS): affiliated with the German Society for Oto-Rhino-Laryngology - Head and Neck Surgery. 2017;274(1):535–42.

18. Stang A. Critical evaluation of the Newcastle-Ottawa scale for the assessment of the quality of nonrandomized studies in meta-analyses. European journal of epidemiology. 2010;25(9):603–5.

19. Hayden JA, van der Windt DA, Cartwright JL, Côté P, Bombardier C. Assessing bias in studies of prognostic factors. Annals of internal medicine. 2013;158(4):280–6.

20. Ikeguchi M. Glasgow prognostic score and neutrophil-lymphocyte ratio are good prognostic indicators after radical neck dissection for advanced squamous cell carcinoma in the hypopharynx. Langenbeck’s archives of surgery. 2016;401(6):861–6.

21. Ong HS, Gokavarapu S, Wang LZ, Tian Z, Zhang CP. Low Pretreatment Lymphocyte-Monocyte Ratio and High Platelet-Lymphocyte Ratio Indicate Poor Cancer Outcome in Early Tongue Cancer.

22. Journal of oral and maxillofacial surgery: official journal of the American Association of Oral and Maxillofacial Surgeons. 2016.

23. Selzer E, Grah A, Heiduschka G, Kornek G, Thurnher D. Primary radiotherapy or postoperative radiotherapy in patients with head and neck cancer: Comparative analysis of inflammation-based prognostic scoring systems. Strahlentherapie und Onkologie: Organ der Deutschen Rontgengesellschaft [et al]. 2015;191(6):486–94.

24. Kano S, Homma A, Hatakeyama H, Mizumachi T, Sakashita T, Kakizaki T, et al. Pretreatment lymphocyte-to-monocyte ratio as an independent prognostic factor for head and neck cancer. Head & neck. 2017;39(2):247–53.

25. Turri-Zanoni M, Salzano G, Lambertoni A, Giovannardi M, Karligkiotis A, Battaglia P, et al. Prognostic value of pretreatment peripheral blood markers in paranasal sinus cancer: Neutrophil-to-lymphocyte and platelet-to-lymphocyte ratio. Head & neck. 2016.

26. Li J-P, Chen S-L, Liu X-M, He X, Xing S, Liu Y-J, et al. A Novel Inflammation-Based Stage (I Stage) Predicts Overall Survival of Patients with Nasopharyngeal Carcinoma. International Journal of Molecular Sciences. 2016;17(11):1900.

27. Duval S, Tweedie R. Trim and fill: a simple funnel□plot–based method of testing and adjusting for publication bias in meta□analysis. Biometrics. 2000;56(2):455–63.

28. Rosenthal R. The file drawer problem and tolerance for null results. Psychological bulletin. 1979;86(3):638.

29. Review Manager (RevMan) [Computer program]. Version 5.3. Copenhagen: The Nordic Cochrane Centre, The Cochrane Collaboration, 2014.

30. Van Rhee, H.J., Suurmond, R., & Hak, T. User manual for Meta-Essentials: Workbooks for meta-analysis (Version 1.0) Rotterdam, The Netherlands: Erasmus Research Institute of Management. 2015. Retrieved from www.erim.eur.nl/research-support/meta-essentials.

31. Koestler DC, Usset JL, Christensen BC, Marsit CJ, Karagas MR, Kelsey KT, et al. DNA methylation-derived neutrophil-to-lymphocyte ratio: an epigenetic tool to explore cancer inflammation and outcomes. Cancer Epidemiology and Prevention Biomarkers. 2016:cebp. 0461.2016.

32. Nakahira M, Sugasawa M, Matsumura S, Kuba K, Ohba S, Hayashi T, et al. Prognostic role of the combination of platelet count and neutrophil–lymphocyte ratio in patients with hypopharyngeal squamous cell carcinoma. European Archives of Oto-Rhino-Laryngology. 2016;273(11):3863–7.

33. Park H-C, Kim M-Y, Kim C-H. C-reactive protein/albumin ratio as prognostic score in oral squamous cell carcinoma. Journal of the Korean Association of Oral and Maxillofacial Surgeons. 2016;42(5):243–50.

34. Acharya S, Rai P, Hallikeri K, Anehosur V, Kale J. Preoperative platelet lymphocyte ratio is superior to neutrophil lymphocyte ratio to be used as predictive marker for lymph node metastasis in oral squamous cell carcinoma. Journal of investigative and clinical dentistry. 2016.

35. An X, Ding P-R, Wang F-H, Jiang W-Q, Li Y-H. Elevated neutrophil to lymphocyte ratio predicts poor prognosis in nasopharyngeal carcinoma. Tumor Biology. 2011;32(2):317–24.

36. Chang H, Gao J, Xu B, Guo S, Lu R, Li G, et al. Haemoglobin, neutrophil to lymphocyte ratio and platelet count improve prognosis prediction of the TNM staging system in nasopharyngeal carcinoma: development and validation in 3237 patients from a single institution. Clinical Oncology. 2013;25(11):639–46.

37. Chen S, Guo J, Feng C, Ke Z, Chen L, Pan Y. The preoperative platelet–lymphocyte ratio versus neutrophil–lymphocyte ratio: which is better as a prognostic factor in oral squamous cell carcinoma? Therapeutic advances in medical oncology. 2016;8(3):160–7.

38. Duzlu M, Karamert R, Tutar H, Karaloglu F, Sahin M, Cevizci R. Neutrophil-lymphocyte ratio findings and larynx carcinoma: a preliminary study in Turkey. Asian Pacific journal of cancer prevention: APJCP. 2014;16(1):351–4.

39. Grimm M, Rieth J, Hoefert S, Krimmel M, Rieth S, Teriete P, et al. Standardized pretreatment inflammatory laboratory markers and calculated ratios in patients with oral squamous cell carcinoma. European Archives of Oto-Rhino-Laryngology. 2016;273(10):3371–84.

40. Jiang R, Zou X, Hu W, Fan Y-Y, Yan Y, Zhang M-X, et al. The elevated pretreatment platelet-to-lymphocyte ratio predicts poor outcome in nasopharyngeal carcinoma patients. Tumor Biology. 2015;36(10):7775–87.

41. Karpathiou G, Giroult J-B, Forest F, Fournel P, Monaya A, Froudarakis M, et al. Clinical and Histologic Predictive Factors of Response to Induction Chemotherapy in Head and Neck Squamous Cell Carcinoma. American Journal of Clinical Pathology. 2016:aqw145.

42. Kum RO, Ozcan M, Baklaci D, Kum NY, Yilmaz YF, Gungor V, et al. Elevated neutrophil-to-lymphocyte ratio in squamous cell carcinoma of larynx compared to benign and precancerous laryngeal lesions. Asian Pacific journal of cancer prevention: APJCP. 2014;15(17):7351–5.

43. Lee C-C, Huang C-Y, Lin Y-S, Chang K-P, Chi C-C, Lin M-Y, et al. Prognostic Performance of a New Staging Category to Improve Discrimination of Disease-Specific Survival in Nonmetastatic Oral Cancer. JAMA Otolaryngology–Head & Neck Surgery. 2017.

44. Li A-C, Xiao W-W, Wang L, Shen G-Z, Xu A-A, Cao Y-Q, et al. Risk factors and prediction-score model for distant metastasis in nasopharyngeal carcinoma treated with intensity-modulated radiotherapy. Tumor Biology. 2015;36(11):8349–57.

45. Li XH, Chang H, Xu BQ, Tao YL, Gao J, Chen C, et al. An inflammatory biomarker□based nomogram to predict prognosis of patients with nasopharyngeal carcinoma: an analysis of a prospective study. Cancer Medicine. 2016.

46. Maruyama Y, Inoue K, Mori K, Gorai K, Shimamoto R, Onitsuka T, et al. Neutrophil-lymphocyte ratio and platelet-lymphocyte ratio as predictors of wound healing failure in head and neck reconstruction. Acta oto-laryngologica. 2017;137(1):106–10.

47. Millrud CR, Mansson Kvarnhammar A, Uddman R, Bjornsson S, Riesbeck K, Cardell LO. The activation pattern of blood leukocytes in head and neck squamous cell carcinoma is correlated to survival. PloS one. 2012;7(12):e51120.

48. Nakashima H, Matsuoka Y, Yoshida R, Nagata M, Hirosue A, Kawahara K, et al. Pre-treatment neutrophil to lymphocyte ratio predicts the chemoradiotherapy outcome and survival in patients with oral squamous cell carcinoma: a retrospective study. BMC cancer. 2016;16(1):41.

49. Ozturk K, Akyildiz NS, Uslu M, Gode S, Uluoz U. The effect of preoperative neutrophil, platelet and lymphocyte counts on local recurrence and survival in early-stage tongue cancer. European archives of oto-rhino-laryngology: official journal of the European Federation of Oto-Rhino-Laryngological Societies (EUFOS): affiliated with the German Society for Oto-Rhino-Laryngology - Head and Neck Surgery. 2016;273(12):4425–9.

50. Perisanidis C, Kornek G, Pöschl PW, Holzinger D, Pirklbauer K, Schopper C, et al. High neutrophil-to-lymphocyte ratio is an independent marker of poor disease-specific survival in patients with oral cancer. Medical Oncology. 2013;30(1):334.

51. Suzuki R, Takagi T, Hikichi T, Konno N, Sugimoto M, Watanabe KO, et al. Derived neutrophil/lymphocyte ratio predicts gemcitabine therapy outcome in unresectable pancreatic cancer. Oncology letters. 2016;11(5):3441–5.

52. Rassouli A, Saliba J, Castano R, Hier M, Zeitouni AG. Systemic inflammatory markers as independent prognosticators of head and neck squamous cell carcinoma. Head & neck. 2015;37(1):103–10.

53. Salim DK, Mutlu H, Eryilmaz MK, Salim O, Musri FY, Tural D, et al. Neutrophil to lymphocyte ratio is an independent prognostic factor in patients with recurrent or metastatic head and neck squamous cell cancer. Molecular and clinical oncology. 2015;3(4):839–42.

54. Trellakis S, Bruderek K, Dumitru CA, Gholaman H, Gu X, Bankfalvi A, et al. Polymorphonuclear granulocytes in human head and neck cancer: enhanced inflammatory activity, modulation by cancer cells and expansion in advanced disease. International journal of cancer. 2011;129(9):2183–93.

55. Tsai YD, Wang CP, Chen CY, Lin LW, Hwang TZ, Lu LF, et al. Pretreatment circulating monocyte count associated with poor prognosis in patients with oral cavity cancer. Head & neck. 2014;36(7):947–53.

56. Valero C, Pardo L, Lopez M, Garcia J, Camacho M, Quer M, et al. Pretreatment count of peripheral neutrophils, monocytes, and lymphocytes as independent prognostic factor in patients with head and neck cancer. Head & neck. 2017;39(2):219–26.

57. Young CA, Murray LJ, Karakaya E, Thygesen HH, Sen M, Prestwich RJ. The prognostic role of the neutrophil-to-lymphocyte ratio in oropharyngeal carcinoma treated with chemoradiotherapy. Clinical Medicine Insights Oncology. 2014;8:81.

58. Yılmaz B, Şengül E, Gül A, Alabalık U, Özkurt FE, Akdağ M, et al. Neutrophil–lymphocyte ratio as a prognostic factor in laryngeal carcinoma. Indian Journal of Otolaryngology and Head & Neck Surgery.1–5.

59. Chen F, Lin L, Yan L, Qiu Y, Cai L, He B. Preoperative Neutrophil-to-Lymphocyte Ratio Predicts the Prognosis of Oral Squamous Cell Carcinoma: A Large-Sample Prospective Study. Journal of Oral and Maxillofacial Surgery. 2016.

60. Donskov F. Immunomonitoring and prognostic relevance of neutrophils in clinical trials. Seminars in cancer biology. 2013;23(3):200–7.

61. Sideras K, Kwekkeboom J. Cancer inflammation and inflammatory biomarkers: can neutrophil, lymphocyte, and platelet counts represent the complexity of the immune system? Transplant International. 2014;27(1):28–31.

62. Wu F, Wu L, Zhu L. Neutrophil to lymphocyte ratio in peripheral blood: a novel independent prognostic factor in patients with head and neck squamous cell carcinoma. Zhonghua zhong liu za zhi [Chinese journal of oncology]. 2017;39(1):29.

63. Zhao GF, Hu YH, Liu RL, Shi F, Li HP, Wang DH, et al. [Clinical significance of the preoperative neutrophil lymphocyte ratio in the evaluation of the prognosis of laryngeal carcinoma]. Zhonghua er bi yan hou tou jing wai ke za zhi = Chinese journal of otorhinolaryngology head and neck surgery. 2016;51(2):112–6.

64. Ahn HK, Hwang IC, Lee JS, Sym SJ, Cho EK, Shin DB. Neutrophil-lymphocyte ratio predicts survival in terminal cancer patients. Journal of palliative medicine. 2016;19(4):437–41.

65. Chua W, Clarke SJ, Charles KA. Systemic inflammation and prediction of chemotherapy outcomes in patients receiving docetaxel for advanced cancer. Supportive Care in Cancer. 2012;20(8):1869–74.

66. Jin Y, Ye X, He C, Zhang B, Zhang Y. Pretreatment neutrophil-to-lymphocyte ratio as predictor of survival for patients with metastatic nasopharyngeal carcinoma. Head & neck. 2015;37(1):69–75.

67. Chen C, Sun P, Dai Q-s, Weng H-w, Li H-p, Ye S. The Glasgow Prognostic Score predicts poor survival in cisplatin-based treated patients with metastatic nasopharyngeal carcinoma. PloS one. 2014;9(11):e112581.

68. Bobdey S, Ganesh B, Mishra P, Jain A. Role of Monocyte Count and Neutrophil-to-Lymphocyte Ratio in Survival of Oral Cancer Patients. International Archives of Otorhinolaryngology. 2017;21(01):21–7.

69. Charles KA, Harris BD, Haddad CR, Clarke SJ, Guminski A, Stevens M, et al. Systemic inflammation is an independent predictive marker of clinical outcomes in mucosal squamous cell carcinoma of the head and neck in oropharyngeal and non-oropharyngeal patients. BMC cancer. 2016;16:124.

70. Chua MLK, Tan SH, Kusumawidjaja G, Shwe MTT, Cheah SL, Fong KW, et al. Neutrophil-to-lymphocyte ratio as a prognostic marker in locally advanced nasopharyngeal carcinoma: A pooled analysis of two randomised controlled trials. European Journal of Cancer. 2016;67:119–29.

71. Fang HY, Huang XY, Chien HT, Chang JT, Liao CT, Huang JJ, et al. Refining the role of preoperative C-reactive protein by neutrophil/lymphocyte ratio in oral cavity squamous cell carcinoma. The Laryngoscope. 2013;123(11):2690–9.

72. Fu Y, Liu W, OuYang D, Yang A, Zhang Q. Preoperative neutrophil-to-lymphocyte ratio predicts long-term survival in patients undergoing total laryngectomy with advanced laryngeal squamous cell carcinoma: a single-center retrospective study. Medicine. 2016;95(6).

73. He JR, Shen GP, Ren ZF, Qin H, Cui C, Zhang Y, et al. Pretreatment levels of peripheral neutrophils and lymphocytes as independent prognostic factors in patients with nasopharyngeal carcinoma. Head & neck. 2012;34(12):1769–76.

74. Kawakita D, Tada Y, Imanishi Y, Beppu S, Tsukahara K, Kano S, et al. Impact of hematological inflammatory markers on clinical outcome in patients with salivary duct carcinoma: A multiinstitutional study in Japan. Oncotarget. 2017;8(1):1083–91.

75. Kim IS, Park SG, Kim H, Choi YJ, Seol YM. Prognostic value of posttreatment neutrophil– lymphocyte ratio in head and neck squamous cell carcinoma treated by chemoradiotherapy. Auris, nasus, larynx. 2017;44(2):199–204.

76. Ma H, Lin Y, Wang L, Rao H, Xu G, He Y, et al. Primary lymphoepithelioma-like carcinoma of salivary gland: sixty-nine cases with long-term follow-up. Head & neck. 2014;36(9):1305–12.

77. Moon H, Roh JL, Lee SW, Kim SB, Choi SH, Nam SY, et al. Prognostic value of nutritional and hematologic markers in head and neck squamous cell carcinoma treated by chemoradiotherapy. Radiotherapy and oncology: journal of the European Society for Therapeutic Radiology and Oncology. 2016;118(2):330–4.

78. Rachidi S, Wallace K, Wrangle JM, Day TA, Alberg AJ, Li Z. Neutrophil-to-lymphocyte ratio and overall survival in all sites of head and neck squamous cell carcinoma. Head & neck. 2016;38 Suppl 1:E1068–74.

79. Rosculet N, Zhou XC, Ha P, Tang M, Levine MA, Neuner G, et al. Neutrophil□to□lymphocyte ratio: Prognostic indicator for head and neck squamous cell carcinoma. Head & neck. 2017.

80. Song Y, Liu H, Gao L, Liu X, Ma L, Lu M, et al. Preoperative neutrophil-to-lymphocyte ratio as prognostic predictor for hypopharyngeal squamous cell carcinoma after radical resections. The Journal of craniofacial surgery. 2015;26(2):e137–40.

81. Sun W, Zhang L, Luo M, Hu G, Mei Q, Liu D, et al. Pretreatment hematologic markers as prognostic factors in patients with nasopharyngeal carcinoma: Neutrophil-lymphocyte ratio and platelet-lymphocyte ratio. Head & neck. 2016;38 Suppl 1:E1332–40.

82. Tu X-P, Qiu Q-H, Chen L-S, Luo X-N, Lu Z-M, Zhang S-Y, et al. Preoperative neutrophil-to-lymphocyte ratio is an independent prognostic marker in patients with laryngeal squamous cell carcinoma. BMC cancer. 2015;15(1):743.

83. Wang J, Wang S, Song X, Zeng W, Wang S, Chen F, et al. The prognostic value of systemic and local inflammation in patients with laryngeal squamous cell carcinoma. OncoTargets and therapy. 2016;9:7177.

84. Wong BY, Stafford ND, Green VL, Greenman J. Prognostic value of the neutrophil-to-lymphocyte ratio in patients with laryngeal squamous cell carcinoma. Head & neck. 2016;38 Suppl 1:E1903–8.

85. Zeng Y-C, Chi F, Xing R, Xue M, Wu L-N, Tang M-Y, et al. Pre-treatment neutrophil-to-lymphocyte ratio predicts prognosis in patients with locoregionally advanced laryngeal carcinoma treated with chemoradiotherapy. Japanese journal of clinical oncology. 2016;46(2):126–31.

86. Amin MB, Edge SB, Greene FL, et al, eds. AJCC Cancer Staging Manual. 8th ed. New York: Springer; 2017.

87. Lu H, Ouyang W, Huang C. Inflammation, a key event in cancer development. Molecular cancer research. 2006;4(4):221–33.

88. Petrie H, Klassen L, Kay H. Inhibition of human cytotoxic T lymphocyte activity in vitro by autologous peripheral blood granulocytes. The Journal of Immunology. 1985;134(1):230–4.

89. El-Hag A, Clark R. Immunosuppression by activated human neutrophils. Dependence on the myeloperoxidase system. The Journal of Immunology. 1987;139(7):2406–13.

90. Gooden MJ, de Bock GH, Leffers N, Daemen T, Nijman HW. The prognostic influence of tumour-infiltrating lymphocytes in cancer: a systematic review with meta-analysis. British journal of cancer. 2011;105(1):93–103.

91. Zhao Q-T, Yang Y, Xu S, Zhang X-P, Wang H-E, Zhang H, et al. Prognostic role of neutrophil to lymphocyte ratio in lung cancers: a meta-analysis including 7,054 patients. OncoTargets and therapy. 2015;8:2731.

92. Li MX, Liu XM, Zhang XF, Zhang JF, Wang WL, Zhu Y, et al. Prognostic role of neutrophil□to□lymphocyte ratio in colorectal cancer: A systematic review and meta□analysis. International journal of cancer. 2014;134(10):2403–13.

93. Hu K, Lou L, Ye J, Zhang S. Prognostic role of the neutrophil–lymphocyte ratio in renal cell carcinoma: a meta-analysis. BMJ open. 2015;5(4):e006404.

94. McMillan DC. The systemic inflammation-based Glasgow Prognostic Score: a decade of experience in patients with cancer. Cancer treatment reviews. 2013;39(5):534–40.

